# Complex enzyme active site scaffolding by iteratively detuned catalytic guidance

**DOI:** 10.64898/2026.10.02.756307

**Authors:** Tingli Qian, Weijin Wang, Shilong Gao, Aiko Muraishi, Beibei Zhao, Davi Nakajima An, Fanglei Xue, Max Kazman, Johnny Mendoza, Hannah Nguyen, Asim Bera, Alex Kang, Avery Yang, Wenzhao Dai, Jia Liang, Yujia Wang, Xiongyi Huang, Frank DiMaio, David Baker

**Affiliations:** Institute for Protein Design, University of Washington, Seattle, WA 98105, USA; Department of Biochemistry, University of Washington, Seattle, WA 98105, USA; Graduate Program in Molecular Engineering, University of Washington, Seattle, WA 98105, USA; Paul G. Allen School of Computer Science and Engineering, University of Washington, Seattle, WA 98105, USA; Department of Chemistry, Johns Hopkins University, Baltimore, MD 21218, USA; Graduate Program in Biological Physics, Structure and Design, University of Washington, Seattle, WA 98105, USA; Graduate Program in Bioengineering, University of Washington, Seattle, WA 98105, USA; Howard Hughes Medical Institute, University of Washington, Seattle, WA 98105, USA

## Abstract

De novo enzyme design requires sequences whose structures reproduce prescribed catalytic geometries. This remains challenging for active sites that couple multiple catalytic residues, cofactors and substrates. Here we introduce the Catalytic Guidance Engine (CaGE), which optimizes designed sequences to satisfy complex sets of functional constraints by alternating constraint guided structure prediction trajectories with sequence design, gradually fading the strength of the constraint information over cycles to nudge the designed sequence towards satisfying all the constraints. We train RF3 to accept atom-pair distances and other design conditions, and use classifier-free guidance to fade their influence over multiple design cycles. We also develop training-free motif guidance protocols which operate at inference time by directly adjusting selected atoms towards the design target geometry during diffusion sampling. CaGE improves catalytic-motif geometry across a diverse enzyme benchmark and yields sequences that recover complex target active sites in independent AF3 predictions. We experimentally demonstrate the power of CaGE by designing active enzymes with complex catalytic sites: a nonheme-iron design positions four active-site residues around an iron centre and a transition-state ligand, and a PLP-dependent design positions four catalytic residues around a PLP cofactor with multiple functional groups. Active site substitutions and a 2.7Å crystal structure support the designed mechanisms and structures. Thus, CaGE provides a route to functional de novo enzymes with complex active sites.

## Main

De novo enzyme design seeks to create protein sequences that fold into stable structures while positioning catalytic functional groups and ligands with atomic precision.^1–5^ Recent deep-learning- based approaches address different parts of this problem, including enzyme-sequence generators such as ProtGPT2 and ZymCTRL, reaction-and-pocket-conditioned models such as EnzymeFlow and GENzyme, and structure-generative motif-scaffolding models including RFdiffusion, RFdiffusion2 (RFD2) and RFdiffusion3 (RFD3)^6–20^. It is not only necessary to generate protein structures that harbor the desired site, but also to identify sequences which are confidently predicted to fold to these structures; a current challenge is that as the complexity of the active site increases, it becomes more and more difficult to identify designs predicted to precisely recapitulate the site. Iterative sequence–structure optimization can improve design qualities,^21,22^ but current approaches provide limited ability to improve catalytic geometry. While structure prediction models have template-based backbone constraints, for enzyme active sites what is critical is the geometry of the active site and in particular the side chain–small molecule interactions, and these are not readily specified in the above methods.

Given an atomic level active site description, and candidate designs harboring the active site, we set out to develop a general method for optimizing both the structure and sequence to achieve high precision active site recapitulation. We reasoned that this optimization could be achieved by iteratively guiding structure prediction calculations toward the designed structure and the specified catalytic geometry, and then progressively removing this guidance, analogous to teacher-policy annealing in reinforcement-learning in which expert supervision supports early learning and is progressively withdrawn as training proceeds.^23,24^ Whereas text-to-image diffusion models commonly use inference-time guidance scales greater than 1 to strengthen conditioning,^25,26^ we reasoned that such constraint fading could be achieved by decreasing inference guidance below 1 across design cycles. We set out to explore this approach which we call CaGE– catalytic guidance engine for de novo design of enzymes with complex active sites.

## Results

Figure 1 outlines the basic idea underlying our approach. While a design model may place catalytic residues and ligands in the intended geometry, structure predictions based on the designed sequence often fail to recover this arrangement (Fig. 1a). This problem is typically overcome by generating large numbers of designs and filtering for those with predicted structures satisfying the input constraints, but for complex constraint sets the fraction of such internally consistent design can be extremely low, making this approach computationally prohibitive. To bring structure predictions closer to the designed geometry, we began with the RosettaFold3 (RF3)^27^ structure prediction method, trained it with design conditions and incorporated guidance. Incorporation of suitably weighted design based active site conditioning as expected brings the generated models to the designed structure (Fig. 1b,c).

**Fig. 1.**
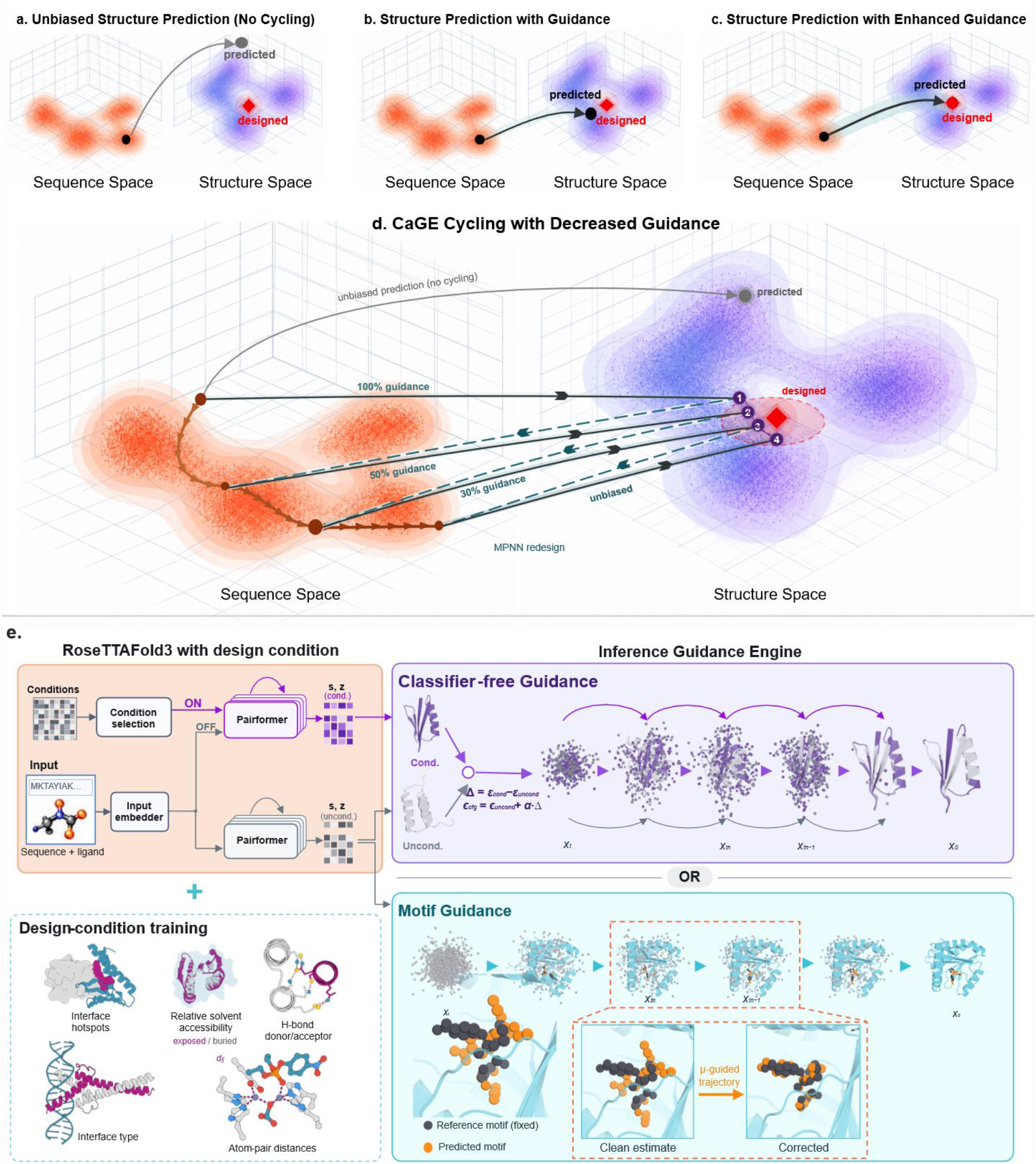
| CaGE combines guidance during iterative enzyme design. **a–c**, Schematic sequence (orange) and structure (purple) landscapes describing sequence-structure relationship. **a**, Without guidance, a candidate sequence (black circle) yields a predicted structure (grey circle) far from the designed target (red diamond); the grey arrow denotes structure prediction. **b**, Guidance biases the predicted structure (black circle) towards the designed structure. **c**, Stronger guidance brings the predicted structure closer to the designed target. **d**, CaGE alternates structure prediction (dark-grey solid arrows, sequence to structure space) with LigandMPNN sequence redesign (teal dashed arrows, structure to sequence space). Guidance decreases across successive design cycles from 100% to 50%, 30% and finally zero. Orange arrows trace changes in sequence space, and numbered purple circles indicate successive predicted structures approaching the target region. A single unguided prediction without cycling (thin grey arrow) remains far from the target. **e,** RF3 is trained to accept design conditions including interface hotspots, relative solvent accessibility, hydrogen-bond donor and acceptor identities, interface type and atom-pair distances. Conditional and unconditional branches share model weights and provide representations for diffusion sampling. CFG adds condition-specific denoising residuals to the unconditional direction, with their contributions controlled by α. Motif guidance directly adjusts selected atoms in the predicted clean structure towards reference coordinates after rigid-group alignment, with correction strength controlled by μ. Corrections are applied at eligible denoising steps, leaving non-motif atoms unchanged by the correction. The inset illustrates the predicted motif (orange) approaching the aligned reference motif (grey). The boxed transition from *x_tn_* to *x_tn_*_−1_ highlights one diffusion update, with the inset showing the motif correction applied to the clean estimate. The two trajectories illustrate CFG and motif guidance as alternative mechanisms for geometric control; the mechanisms can also be combined, as described in Methods.

However, structure–sequence agreement under guidance does not ensure that the sequence will recover the site when this guidance is removed. To overcome this problem, we turned to classifier free guidance,^25^ which combines conditional and unconditional denoising directions at adjustable scales (see below and Methods). Whereas inference-time guidance in image-generation diffusion models commonly uses scales greater than 1 to strengthen conditioning, we reasoned that iterative guidance with scales lower than 1 could be used to progressively withdraw guidance over successive design cycles to arrive at designed sequences which encode the designed sites. (Fig. 1d).

### Incorporation of constraint conditioning in RF3

CAGE requires the ability to condition a structure prediction method on constraint sets similar to those used in design calculations. While structure prediction methods such as AF3 or Boltz,^28,29^ allow template-based backbone constraints, for enzyme active sites what is critical is the geometry of the active site and in particular the side chain–small molecule interactions, and these are not readily specified in the above methods.

We extended RF3 training to incorporate additional design conditions. RF3 has an overall AF3- like architecture, with a pairformer trunk feeding a diffusion module that generates all-atom coordinates. We extended its training to incorporate five types of design condition: interface hotspots, relative solvent accessibility, hydrogen-bond donor and acceptor identities, interface type, and atom-pair distances (Fig. 1e and Supplementary Fig. 1). Interface hotspots inform target binding sites (Supplementary Figure 2), whereas relative solvent accessibility specifies the desired exposure or burial of selected atoms to water. Hydrogen-bond conditions identify atoms intended to act as donors or acceptors, and interface-type conditioning specifies the class of molecular interface, such as a protein–DNA interface. Atom-pair distance conditioning specifies the identities of two atoms and their target separation without imposing an absolute coordinate frame.

### Classifier-free guidance combines and adjusts the strength of design conditions

With these design conditions learned by the structure prediction model, we controlled the relative influence of the different design conditions on the final structure output with classifier-free guidance. Using CFG, multiple design conditions can be combined at independently adjustable strengths, allowing their contributions to be independently and progressively reduced across design cycles. For each condition *i*, the CFG scale α*ᵢ* interpolates between the unconditional prediction (α*ᵢ* = 0) and the conditional prediction (α*ᵢ* = 1): setting α*ᵢ* = 0 removes its contribution entirely, whereas α*ᵢ* = 1 retains it at full strength.

For enzyme design, we combined atom-pair distance conditioning with template-based conditioning to guide backbone geometry and ligand placement towards the target theozyme configuration, including catalytic side-chain geometry at atomic resolution^5^. We selected eight enzyme targets spanning diverse catalytic motifs and ligand environments to evaluate whether CaGE could accommodate different geometric constraints on catalytic residues and ligand placement. These comprised purine nucleoside phosphorylase (PNP), ornithine decarboxylase (ODC), 4-chlorobenzoyl-CoA dehalogenase (4-CBA-CoA DH), enoyl-CoA hydratase (crotonase) and citrate synthase (CS) (Fig. 2a), together with UDP-galactose 4-epimerase (GalE), glycosidase (GLY) and UDP-glucose 6-dehydrogenase (UGDH) (Supplementary Fig. 4a). Increasing the classifier-free guidance scale α for atom-pair distance conditioning from 0 to 1 reduced the median motif r.m.s.d. across the tested targets (Figure 2a and Supplementary Figure 4a). At matched CFG scales, RF3 trained with atom-pair distance conditioning achieved lower median motif r.m.s.d. than RF3 without this conditioning (Figure 2a and Supplementary Figure 4a).

**Fig. 2.**
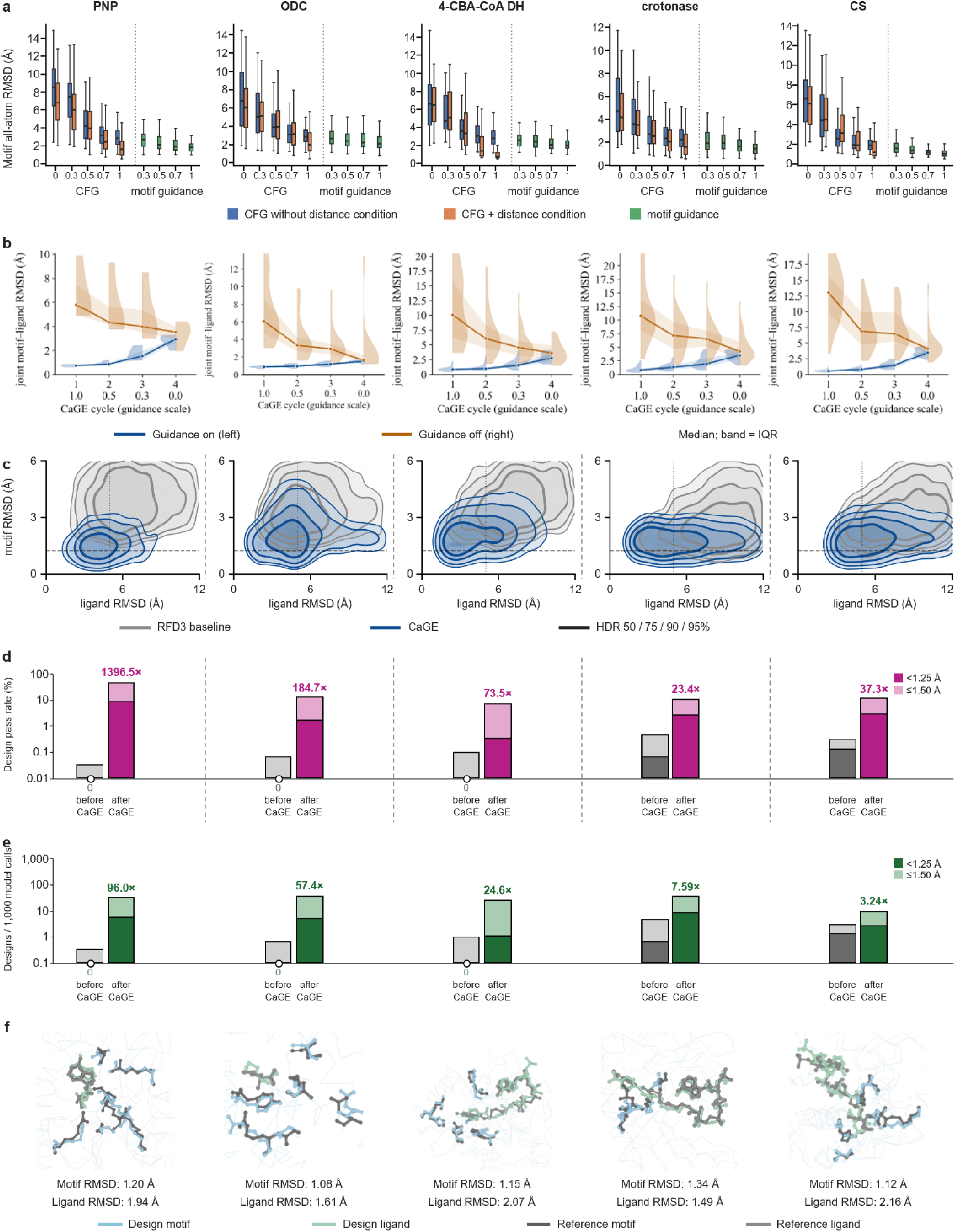
| Structure-conditioned cycling yields catalytic-motif and ligand recovery at the unguided endpoint. a, Guidance steers predictions towards the target geometry. CFG strength α was varied from 0 to 1 without motif guidance, before (blue) and after (orange) atom-pair distance-conditioning training. Motif guidance strength μ was varied from 0 to 1 with α = 1 and without distance- conditioning training (green). Box plots show motif r.m.s.d. under catalytic-backbone alignment. Boxes span the 25th–75th percentiles; horizontal lines indicate medians. **b, Guided and unguided predictions converge across CaGE cycles.** Lineage-tracked sequences were evaluated by cycling predictions and independent unguided refolds using identical sequences through cycles. CFG α and MG μ decreased together through 1.0, 0.5, 0.3 and 0 across cycles 1–4; both were zero in independent refolds. Lines and bands show median joint motif–ligand r.m.s.d. and interquartile ranges. Split violins show cycling predictions (blue, left) and unguided refolds (orange, right). **c, CaGE improves active-site geometry in AF3 predictions.** Joint distributions of sequence-level motif and ligand r.m.s.d. are shown for CaGE (blue) and the RFD3–LigandMPNN baseline (grey). Contours enclose 50%, 75%, 90% and 95% of the probability mass in highest-density regions after normalization within the plotted window. **d, CaGE increases AF3 endpoint pass rates.** The dark portion of each bar shows the pass rate for motif r.m.s.d. <1.25 Å and ligand r.m.s.d. <5 Å; the entire bar shows the pass rate for motif r.m.s.d. ≤1.5 Å and ligand r.m.s.d. ≤5 Å. CaGE is magenta and the baseline grey. Labels indicate CaGE-to-baseline pass-rate ratios at the 1.5-Å motif threshold; zeros are marked near the x axis. **e, CaGE maintains design efficiency despite additional prediction calls.** Efficiency is the number of passing designs per 1,000 RF3 and AF3 prediction calls for CaGE (dark green) and the baseline (grey). Thresholds and bar shading are as in d. Labels indicate CaGE-to-baseline efficiency ratios at the 1.5-Å motif threshold; zeros are marked near the x axis. Full counts are provided in Supplementary Table 3. **f, Structure comparison of AF3 predictions and target theozymes.** Representative AF3 predictions are superposed on target theozymes by alignment of motif Cα atoms. Motif and ligand r.m.s.d. values are indicated. Predicted backbones and catalytic side chains are pale blue, and predicted ligands are green with heteroatoms coloured by element. Reference theozymes and ligands are grey.

### Motif guidance enables precise geometric control without additional training

Training can incorporate a broad range of design conditions, enabling control over diverse molecular properties. To enable utilization of a broader range of structure prediction models, we also developed a training-free alternative for geometric control termed motif guidance. At each denoising step, motif guidance corrects the clean estimate x^₀, thereby steering the diffusion trajectory toward the designed geometry, with μ adjusting the strength of the correction.

Without atom-pair distance-conditioning, increasing motif guidance strength μ from 0 to 1 while holding CFG at its maximum tested scale (α = 1) further reduced motif r.m.s.d. across targets relative to CFG alone (Fig. 2a). On our enzyme-design benchmarks, motif guidance combined with CFG achieved motif accuracy comparable to that obtained with atom-pair distance- conditioning training and CFG. Representative structural comparisons across all targets illustrate progressive improvements in catalytic-motif geometry with increasing guidance strength (Supplementary Fig. 3).

### Guided and unguided predictions converge across cycles

To test the mechanism proposed in Fig. 1d, in which successive rounds of structure prediction and sequence redesign reduce reliance on external guidance as sequences become more compatible with the target active-site geometry, we compared guided predictions with independent unguided refolds of identical amino acid sequences within each cycle. This comparison assessed whether the target active-site geometry could increasingly be recovered without external guidance during iterative prediction and sequence redesign. As guidance progressively decreased, the difference between median joint motif–ligand r.m.s.d. values narrowed monotonically across all eight targets (Fig. 2b and Supplementary Fig. 4b). This convergence supports the proposed transition from guidance-dependent geometry towards geometry that is increasingly supported by the redesigned sequences.

### CaGE recovers active-site geometry in unguided RF3 and AF3 predictions

We next evaluated CaGE across eight enzyme targets, using sequences designed on RFD3- generated backbones without CaGE cycling as the baseline. The final CaGE cycle was completed without guidance, and the final sequences were subsequently evaluated with AF3. Across the five targets shown in Fig. 2, CaGE AF3 predictions had lower median motif and ligand r.m.s.d. values than baseline predictions (Fig. 2c).

We assessed two joint pass criteria: motif r.m.s.d. <1.25 Å and ligand r.m.s.d. <5 Å (strict), or motif r.m.s.d. ≤1.5 Å and ligand r.m.s.d. ≤5 Å (relaxed). Fig. 2d shows the sequence-level pass rate: the percentage of evaluated designed sequences meeting each joint criterion. For the five targets shown in Fig. 2, CaGE increased this pass rate at both motif thresholds. At the 1.5-Å motif r.m.s.d. threshold, CaGE increased pass rates across targets. More notably, at the more stringent 1.25-Å threshold, CaGE recovered passing designs for challenging targets where the baseline yielded none, achieving pass rates of 8.11%, 1.57% and 0.32% for PNP, ODC and 4-CBA-CoA DH, respectively (Fig. 2d). After normalizing with the number of structure-prediction calls for efficiency analysis, CaGE produced more passing sequences per 1,000 prediction calls than the RFD3 baseline for all five targets at both motif cutoffs (Fig. 2e). Among designs passing the 1.5- Å motif criterion, CaGE designs occupied structure clusters not represented among the sampled baseline successes (Supplementary Fig. 5).

At the 1.25-Å motif cutoff, this yield increased from 0.62 to 8.06 for crotonase and from 1.25 to 2.46 for CS. The baseline yielded no passing sequences for PNP, ODC or 4-CBA-CoA DH at this cutoff, whereas CaGE yielded 5.579, 4.889 and 1.078 per 1,000 structure prediction calls, respectively. The advantage persisted for all five targets when the motif cutoff was relaxed to 1.5 Å. Results for the three additional targets, GalE, GLY and UGDH, are reported in Supplementary Fig. 4d,e. Representative AF3 predictions illustrate the recovered catalytic-motif geometries and ligand poses relative to the target theozymes (Fig. 2f).

### CaGE refines a coupled nonheme-iron catalytic motif

To test whether CaGE could recover coupled metal-coordination and reaction-state geometry, we designed scaffolds around a nonheme-iron transition-state motif for a C–N bond-forming rearrangement (Fig. 3a). The motif comprises a mononuclear iron centre, its first-shell coordinating residues and a transition-state ligand spanning the reacting fragments, requiring simultaneous recovery of the coordination sphere and reactive geometry. This motif is challenging to scaffold and recover in structure prediction because it combines a non-native transition-state ligand with multiple coupled degrees of freedom governing metal coordination and the relative positioning of the reacting fragments.

**Fig. 3.**
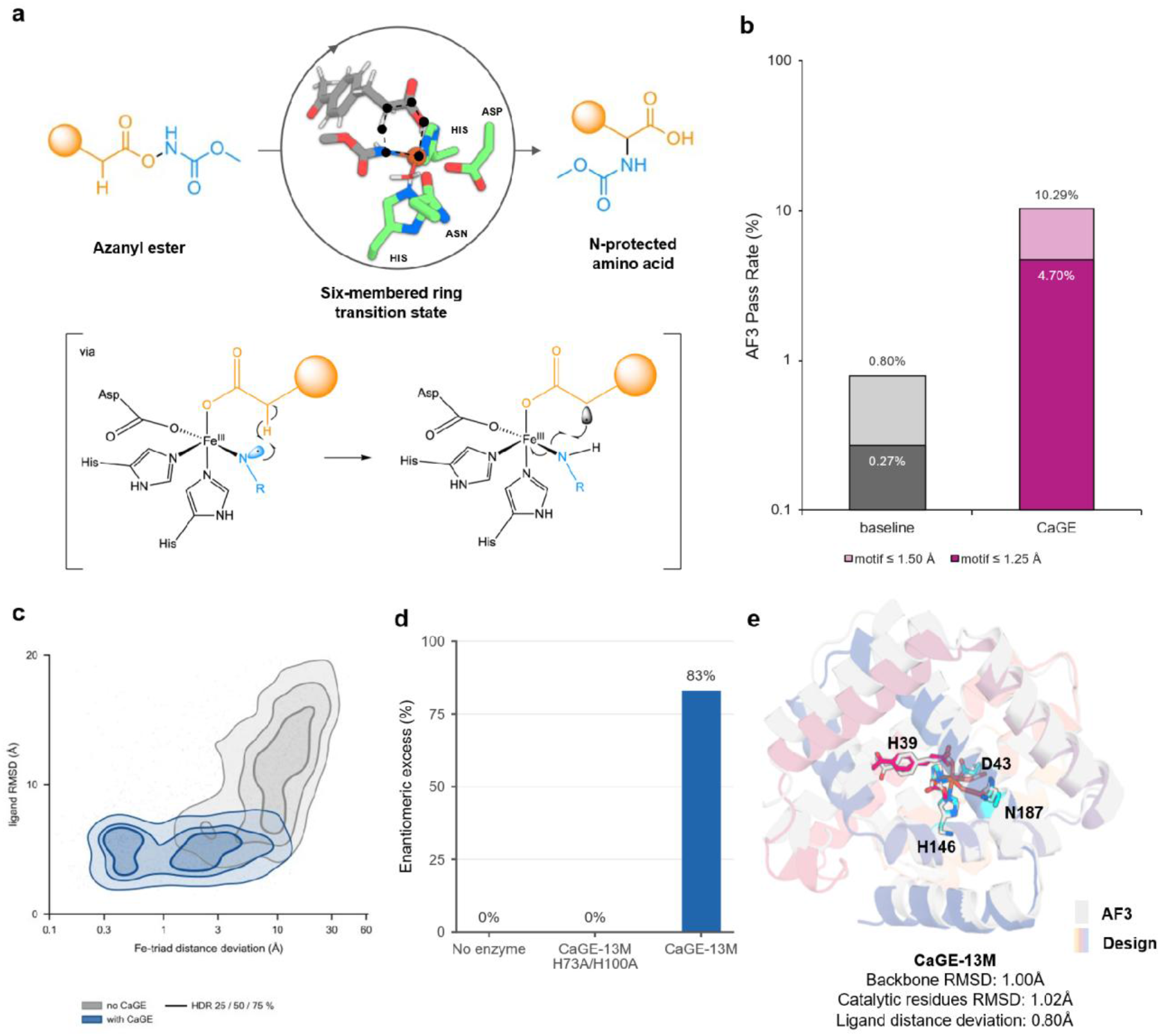
| CaGE refines a coupled nonheme-iron transition-state motif. a, Catalytic motif used for de novo scaffolding, comprising a mononuclear iron centre, its first- shell coordinating residues and a transition-state ligand spanning the reacting fragments. The motif specifies a partially formed bond in addition to the metal-coordination geometry. **b,** Sequence- level recovery before and after one CaGE cycle, starting from 1,500 unoptimised RFdiffusion3– LigandMPNN backbones. After identical duplicate and low-complexity filtering, the baseline comprised 1,491 sequences and the CaGE arm comprised 175 retained parent sequences plus 1,526 redesigns (n = 1,701). Both arms used identical AF3 settings, references and catalytic-residue lists. Passing required mean motif r.m.s.d. ≤ 1.25 Å (dark bars) or ≤ 1.5 Å (total bar heights) and mean ligand r.m.s.d. ≤ 5 Å across five diffusion samples, rather than the sample-wise voting criterion used elsewhere. Motif r.m.s.d. was calculated over catalytic-residue heavy atoms after alignment on their backbone N, CA and C atoms. Rates are normalised to each folded sequence set, excluding backbones discarded during cycling. **c,** Joint distribution of ligand r.m.s.d. and Fe-triad distance deviation for the same two arms. Contours are highest-density regions containing 25, 50 and 75% of the in-window probability mass; the x axis is logarithmic and the density is renormalised within the plotted window (94.8% of baseline and 99.7% of CaGE designs fall inside it). Medians: 8.80 Å triad deviation and 11.40 Å ligand r.m.s.d. without cycling, versus 1.81 Å and 5.39 Å after one cycle. **d**, Product stereoselectivity for **CaGE-13M** relative to a no-enzyme control and the H39A/H147A variant. Enantiomeric analysis yielded 83% ee for **CaGE-13M**, whereas the no- enzyme and mutant reactions were racemic (0% ee). The stereochemical divergence from the racemic control is consistent with chiral control by the designed active site. **e**, Structural models of the first-round hit **CaGE-13M**, from the production design campaign. Design models are superposed with independent AF3 predictions. The active site contains the iron-coordinating residues His39, His147 and Asp43, together with Asn187. Backbone and catalytic-residue r.m.s.d. values are 1.00 and 1.02 Å, respectively; the ligand distance deviation is 0.80 Å.

We compared unoptimised RFdiffusion3–LigandMPNN^30^ designs with sequences generated after one CaGE cycle, starting from the same pool of 1,500 backbones sampled before additional filtering or refinement. Following identical sequence-quality filtering, 1,491 baseline sequences and 1,701 CaGE sequences were evaluated using the same unguided AF3 protocol, with five diffusion samples per sequence. One CaGE cycle increased the fraction meeting a catalytic-motif r.m.s.d. threshold of 1.25 Å from 0.27% to 4.70%, and a threshold of 1.5 Å from 0.80% to 10.29%, with both criteria additionally requiring ligand r.m.s.d. ≤ 5 Å (Fig. 3b). These rates measure recovery per evaluated sequence and exclude backbones discarded during cycling. The broader structural distributions also shifted toward the target geometry: median Fe-triad distance deviation decreased from 8.80 to 1.81 Å, and median ligand r.m.s.d. decreased from 11.40 to 5.39 Å (Fig. 3c). Thus, one CaGE cycle generated sequences whose unguided predictions better recovered both metal coordination and ligand placement.

We next scaled up the CaGE calculations to generate designs to experimentally test. We applied four CaGE refinement cycles, with the CFG scale reduced from 1.0 to zero. Final unguided AF3 evaluation identified 160 designs that satisfied the fold and complete motif criteria in at least three of five diffusion samples. Of 71 designs screened experimentally, six produced product signals above the no-enzyme control. The design CaGE-13M showed activity and yielded products with 83% enantiomeric excess (Fig. 3d). This result supports involvement of the designed active site in controlling the reaction outcome. Together, these results connect improved computational recovery of a coupled catalytic motif with experimental activity and stereoselectivity in a designed nonheme-iron scaffold.

CaGE-13M is a 209-residue protein with a mononuclear iron site coordinated by His39, His147 and Asp43, with Asn187 also positioned in the active site (Fig. 3e). An independent AF3 prediction recovered the backbone and catalytic residues with r.m.s.d. values of 1.00 and 1.02 Å, respectively, and a ligand distance deviation of 0.80 Å.

### De novo design and experimental validation of a PLP-dependent retro-aldolase

Pyridoxal phosphate (PLP) is a versatile biological cofactor that supports diverse transformations, including transamination, decarboxylation, racemization, carbon–carbon bond cleavage and formation, and β- and γ-elimination reactions.^31^ Its reactivity arises from formation of a Schiff base with a catalytic lysine or amino-group-containing substrate, allowing the conjugated pyridinium system to stabilize high-energy carbanionic intermediates.

PLP-dependent catalysis therefore provides a stringent test for de novo enzyme design. Productive catalysis requires the protein to position a catalytic lysine for internal-aldimine formation, recognize the PLP phosphate, tune the electronic state of the pyridine ring, orient the substrate relative to the cofactor and stabilize multiple intermediates along the reaction coordinate. These requirements go substantially beyond simple cofactor binding. To our knowledge, no PLP-dependent enzyme has previously been generated by de novo scaffolding of a predefined PLP catalytic motif.

We therefore applied CaGE to the design of a PLP-dependent retro-aldolase that cleaves a β- hydroxy amino acid into the corresponding aldehyde and glycine through a PLP external aldimine (Fig. 4a). The catalytic motif contained four residues surrounding the PLP–substrate complex (Fig. 4b). Lys142 was positioned to form the resting-state PLP internal aldimine, Asp27 to interact with the PLP pyridine nitrogen, His31 to form a π interaction with the PLP pyridine ring and contact the substrate β-hydroxyl group, and Arg159 to recognize and orient the substrate carboxylate. Together, these interactions defined an atomically specified catalytic arrangement coupling cofactor activation with substrate positioning.

**Fig. 4.**
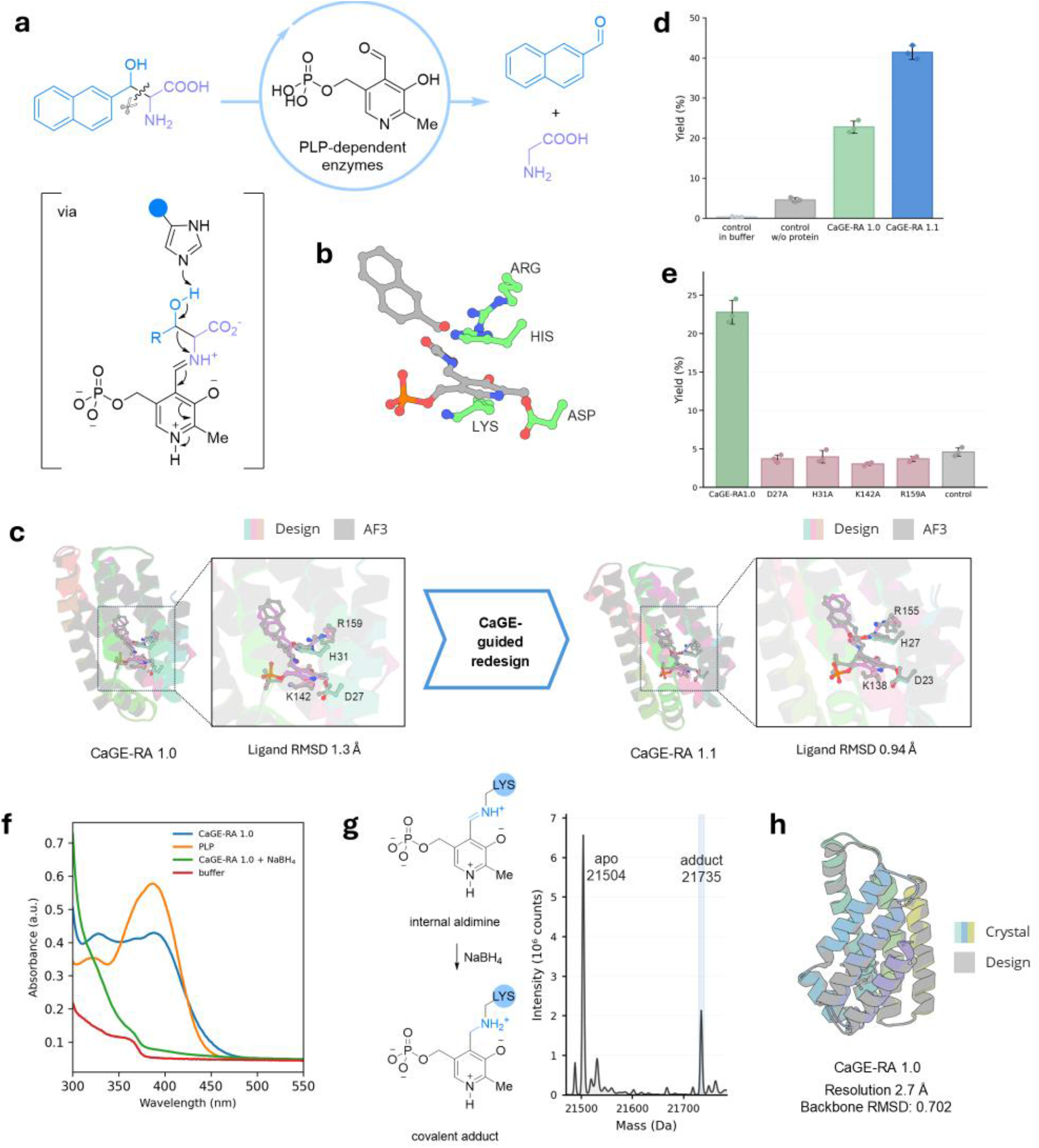
| De novo design and experimental validation of a monomeric PLP-dependent retro-aldolase. a, Target PLP-dependent retro-aldol reaction. Cleavage of the β-hydroxy amino acid substrate affords the corresponding aldehyde and glycine through a PLP external-aldimine intermediate. The bracketed structure illustrates the transition state used to define the catalytic geometry for design. **b,** Catalytic motif used for de novo scaffolding, comprising PLP, the substrate and four catalytic residues. Lys142 is positioned to form the PLP internal aldimine, Asp27 interacts with the PLP pyridine nitrogen, His31 is positioned to form a π interaction with the PLP pyridine ring and to contact the substrate β-hydroxyl group, and Arg159 engages the substrate carboxylate, thereby defining the geometry required for cofactor activation and substrate positioning.**c,** Structural models of the first-round hit **CaGE-RA 1.0** and the second-round redesign **CaGE-RA 1.1**. Design models are superposed with independent AF3 predictions, with enlarged views showing recovery of the designed active-site geometry. CaGE-guided redesign of the experimentally active first-round scaffold yielded **CaGE-RA1.1**. Catalytic-residue r.m.s.d. values were calculated after backbone alignment over the heavy atoms of Asp27, His31, Lys142 and Arg159 in CaGE-RA1.0 and the corresponding residues Asp23, His27, Lys138 and Arg155 in CaGE-RA1.1. **d,** Retro-aldol activity of first- and second-round designs under the screening conditions. Reactions contained 25 μM enzyme, 2.5 mM β-hydroxy amino-acid substrate and 25 μM PLP in 100 mM KPi, pH 8.0, and were incubated at 30 °C for 16 h. Bars show mean ± s.d.; individual measurements are shown as points; n = 3 independent reactions. **e,** Functional interrogation of the designed catalytic motif in **CaGE-RA1.0**. SEC-purified D27A, H31A, K142A and R159A variants were assayed under the same conditions as the parent design (25 μM enzyme, 2.5 mM substrate, 25 μM PLP, 100 mM KPi, pH 8.0, 30 °C, 16 h). All four substitutions reduced product formation to approximately the no-protein background, supporting the functional contribution of the computationally specified residues. Bars show mean ± s.d.; individual measurements are shown as points; n = 3 independent reactions. **f,** UV–visible characterization of PLP association with **CaGE-RA1.0**. Spectra were recorded from 250 to 550 nm for 50 μM **CaGE- RA1.0** with 50 μM PLP in 100 mM KPi, pH 8.0, together with free PLP and buffer controls. Treatment of the **CaGE-RA1.0–PLP** sample with 8 mM NaBH₄ for 20 min on ice strongly suppressed the visible PLP absorption feature, consistent with reduction of a protein-associated Schiff-base species. **g,** Reductive trapping of the PLP internal aldimine. NaBH₄ converts the reversible Lys–PLP Schiff base into a stable covalent adduct. Intact-protein mass spectrometry of NaBH₄-treated **CaGE-RA1.0** showed the apo protein at 21,504 Da and an additional species at 21,735 Da. The 231-Da mass increase is consistent with incorporation of a single PLP molecule following Schiff-base formation and reduction. **h,** Crystal structure of apo **CaGE-RA1.0**. The 2.7- Å crystal structure superposes closely with the design model, with a backbone r.m.s.d. of 0.702 Å, confirming recovery of the intended de novo fold. **CaGE-RA1.0** forms a compact monomeric architecture, demonstrating that a functional PLP active site can be constructed within a single designed chain rather than relying on the oligomeric architectures used by canonical natural PLP- dependent aldolases.

Starting from this motif, candidate scaffolds were generated using RFdiffusion3, LigandMPNN and iterative FastRelax^32^–LigandMPNN refinement before CaGE cycling. None of the 30,293 pre- CaGE sequences passed the final AF3 geometric criteria when evaluated under the same protocol used for the CaGE candidates. CaGE refinement yielded 1,029 variants passing the final unguided RF3 filters, of which 96 also passed independent AF3 validation without a protein template or supplied ligand pose.

A lysate-based primary screen of the 96 computationally selected first-round designs identified **CaGE-RA1.0** as an active hit (Supplementary Fig. 7). Primary screening was performed with 2.5 mM β-hydroxy amino-acid substrate and 25 μM PLP in 100 mM potassium phosphate (KPi), pH 8.0, at 30 °C for 16 h. **CaGE-RA1.0** was subsequently expressed at larger scale, purified by Ni-NTA affinity chromatography and size-exclusion chromatography, and its activity was confirmed using purified protein under matched reaction conditions. Its AF3 prediction closely reproduced the design model, with PLP–substrate ligand r.m.s.d. value of 1.30 Å, respectively (Fig. 4c). Thus, the designed geometry was independently recovered from sequence rather than maintained only by structural guidance.

**CaGE-RA1.0** was designed as a compact monomer, with the PLP-binding and catalytic environment contained within a single polypeptide chain. This architecture differs from canonical natural PLP-dependent threonine aldolases,^33^ which typically adopt evolved PLP-enzyme folds relying on oligomeric assemblies. **CaGE-RA1.0** therefore represents a structurally distinct solution in which PLP-dependent carbon–carbon bond chemistry is embedded within a de novo monomeric scaffold.

### Hit-guided redesign improves catalytic activity

We next asked whether an experimentally validated first-round hit could provide a starting point for further CaGE optimization. **CaGE-RA1.0** was used as the structural template for a second design round while retaining the catalytic motif (Fig. 4c). Rather than sampling mutations independently, the redesign allowed coordinated sequence and structural changes around the active site while preserving the validated global architecture. The resulting design, **CaGE-RA1.1**, again showed close agreement between the design model and AF3 prediction, with ligand r.m.s.d. value of 0.94 Å, respectively (Fig. 4c). This redesign also improved catalytic performance. Under matched conditions, **CaGE-RA1.0** produced 22.8 ± 1.6% product, whereas **CaGE-RA1.1** reached 41.4 ± 1.7% yield (mean ± s.d., n = 3 independent reactions), corresponding to an approximately 1.8-fold improvement (Fig. 4d). Thus, the same framework used to generate the initial active scaffold could subsequently improve an experimentally validated hit without a large combinatorial mutagenesis campaign.

### Mechanistic and structural validation of the designed PLP enzyme

To test whether activity depended on the intended catalytic architecture, each of the four designed catalytic residues in **CaGE-RA1.0** was individually replaced with alanine (Fig. 4e). D27A, H31A, K142A and R159A all reduced product formation to approximately the no-protein background, indicating that activity depends on the designed residue network rather than nonspecific PLP-mediated chemistry. The strong effect of K142A is consistent with its proposed role as the covalent anchor for the resting-state PLP internal aldimine. The effects of D27A, H31A and R159A further support the importance of PLP activation and substrate positioning. Thus, residues specified geometrically during computational design were also required experimentally for productive catalysis.

We next examined the chemical state of PLP associated with **CaGE-RA1.0**. UV–visible spectra were recorded for 50 μM purified **CaGE-RA1.0** before and after addition of one molar equivalent of PLP (50 μM) in 100 mM KPi, pH 8.0. The **CaGE-RA1.0–PLP** sample showed a distinct absorption profile compared with free PLP in solution (Fig. 4f), consistent with incorporation of the cofactor into a different electronic environment. Treatment of the **CaGE-RA1.0–PLP** complex with NaBH₄ strongly diminished the visible PLP absorption feature, as expected upon reduction of a protein-bound Schiff base. We therefore used NaBH₄ to trap the proposed Lys142– PLP internal aldimine and analysed the resulting protein by intact mass spectrometry (Fig. 4g). **CaGE-RA1.0** showed a major apo species at 21,504 Da and an additional species at 21,735 Da following PLP loading and reductive trapping. The 231-Da mass increase is consistent with incorporation of a single PLP molecule through Schiff-base formation followed by reduction.

Together with the spectroscopic measurements, these data support formation of the designed covalent PLP–lysine species rather than simple non-covalent cofactor association.

Finally, we determined the crystal structure of **CaGE-RA1.0** at 2.7 Å resolution (Fig. 4h). The experimental structure closely reproduced the computational model, with a backbone r.m.s.d. of 0.702 Å, confirming that the designed global architecture was encoded by the sequence and realized experimentally. **CaGE-RA1.0** also retained the intended monomeric architecture in the crystal structure. This distinguishes the design from canonical PLP-dependent threonine aldolases, in which oligomerization is a common structural feature, and demonstrates that the requirements for PLP-dependent carbon–carbon bond cleavage can be accommodated within a compact single-chain scaffold.

## Discussion

Here, we introduce CaGE, an iterative enzyme-design framework that combines guided structure prediction with sequence redesign and progressively withdraws the guidance to obtain sequences which encode structures satisfying complex sets of constraints. Sequence design methods such as ProteinMPNN find sequences which maximize P(sequence| structure), but the maximum of P(structure|sequence) for these sequences may not be the designed structure; CaGE effectively nudges the designed sequence towards encoding the designed structure. CaGE uses model forward passes rather than backpropagation and so is considerably faster than hallucination approaches which also optimize sequences to encode desired structures^34,35^. Protein Hunter and HalluDesign have shown the effectiveness of iterative prediction-and-redesign cycles for binder design, where protein-protein interactions can be viewed as an inherently known design condition in structure prediction models. CaGE goes considerably beyond these approaches by incorporating a broad range of conditioning information on atom-pair distance distances and other protein properties relevant to enzymatic and other functions, and gradually fading this information during design iterations to nudge the designed sequence to accurately encode the designed constraints. The wide range of conditioning gives CaGE broad versatility, for example, hotspot conditioning could direct antibody binding to selected hotspots on an antigen, interface-type conditioning could specify the class of binding partner for design of multicomponent protein–nucleic acid assemblies, and RASA conditioning could guide ligand and pH-dependent binding proteins.

Our experimental tests show that CaGE can yield functional designs under different catalytic constraints. In the nonheme-iron system, **CaGE-13M** produced a product signal above background with 83% enantiomeric excess, whereas the no-enzyme and coordination-site-mutant controls were racemic. The PLP-dependent retro-aldolase has a crystal structure nearly identical to the design model, substitutions at the four designed catalytic positions reduce product formation to near background, and spectroscopy and reductive trapping support formation of the intended protein– PLP Schiff-base species. CaGE refinement increased the endpoint product yield, illustrating a further use of the workflow in improving an experimentally identified hit.

CaGE illustrates how a structure predictor can act as a design engine, guiding iterative changes to sequences and structures toward a prescribed catalytic geometry. Our inference-time guidance strategies enable use of multiple off the shelf prediction methods in CaGE, and could further improve performance by avoiding possible biases associated with a single method. More broadly, CaGE shows how learned sequence–structure relationships can support both prediction and controlled molecular design, suggesting a foundation for biological world models that predict the consequences of molecular changes and guide interventions toward desired functions.

## Acknowledgements

We thank Luki Goldschmidt and the IT team, and Kandise VanWormer and the lab management team, for maintaining the computational and wet-lab resources at the Institute for Protein Design. We thank the RF3 development team for developing the underlying structure-prediction model. We also thank Andrew Kubaney for proofreading the manuscript and for helpful discussions and suggestions during its revision. X-ray diffraction data were collected at the Frontier Microfocusing Macromolecular Crystallography (FMX) beamline 17-ID-2 of the National Synchrotron Light Source II (NSLS-II). The Center for BioMolecular Structure (CBMS) is primarily supported by the National Institutes of Health, National Institute of General Medical Sciences (NIGMS) through a Center Core P30 Grant (P30GM133893), and by the US Department of Energy Office of Biological and Environmental Research (KP1605010).

## Author contributions

Research design: T.Q., D.B.; Development of design-condition training in RF3: T.Q., D.N.A., F.D., A.M., M.K.,; Implementation of classifier-free guidance and motif guidance: T.Q.,; Benchmark: T.Q.; Nonheme-iron design campaign: S.G.; Synthesis of substrates and product standards, and chiral LC–MS analysis of nonheme-iron designs: B.Z., S.G., X.H.; PLP retro-aldolase design campaign: W.W., Y.W., W.D.; Gene synthesis, protein expression, purification and activity assays: W.W.; UV–visible spectroscopy, NaBH₄ trapping and intact-protein mass spectrometry: W.W.; Crystallization, data collection and structure determination: J.M., H.N., A.B., A.K.; Infrastructure and project set-up: F.D., F.X., A.Y., J.L., Y.W.; Figure creation: T.Q., W.W., S.G.; Writing — original draft: T.Q., W.W., S.G., D.B.; Review and editing: all authors; Supervision: D.B., F.D., X.H.

## Materials and correspondence

Correspondence and requests for materials should be addressed to David Baker.

## Data availability

The data supporting this study are provided in the Supplementary Information and accompanying Source Data files. Atomic coordinates and structure factors for apo CaGE-RA1.0 will be available from the Protein Data Bank under accession code [PDB ID; accession code will be updated upon acceptance].

## Code availability

Code for running CaGE is available under the MIT open-source license on GitHub via https://github.com/baker-laboratory/CaGE.

## References

1 Bornscheuer, U. T. et al. Engineering the third wave of biocatalysis. Nature 485, 185–194 (2012). 10.1038/nature11117

2 Buller, R. et al. From nature to industry: Harnessing enzymes for biocatalysis. Science 382, eadh8615 (2023). 10.1126/science.adh8615

3. 3 Arnold, F. H. Directed Evolution: Bringing New Chemistry to Life. Angew. Chem. Int. Ed. 57, 4143–4148 (2018). 10.1002/anie.201708408

4 Turner, N. J. Directed evolution drives the next generation of biocatalysts. Nat. Chem. Biol. 5, 567– 573 (2009). 10.1038/nchembio.203

5 Lovelock, S. L. et al. The road to fully programmable protein catalysis. Nature 606, 49–58 (2022). 10.1038/s41586-022-04456-z

6 Ferruz, N., Schmidt, S. & Höcker, B. ProtGPT2 is a deep unsupervised language model for protein design. Nat. Commun. 13, 4348 (2022). 10.1038/s41467-022-32007-7

7 Munsamy, G. et al. Conditional language models enable the efficient design of proficient enzymes. bioRxiv (2024). 10.1101/2024.05.03.592223

8 Hua, C. et al. EnzymeFlow: Generating reaction-specific enzyme catalytic pockets through flow matching and co-evolutionary dynamics. arXiv preprint arXiv:2410.00327 (2024). 10.48550/arXiv.2410.00327

9 Hua, C. et al. Reaction-conditioned de novo enzyme design with GENzyme. arXiv preprint arXiv:2411.16694 (2024). 10.48550/arXiv.2411.16694

10 Watson, J. L. et al. De novo design of protein structure and function with RFdiffusion. Nature 620, 1089–1100 (2023). 10.1038/s41586-023-06415-8

11 Braun, M. et al. Computational enzyme design by catalytic motif scaffolding. Nature 649, 237–245 (2026). 10.1038/s41586-025-09747-9

12 Jiang, L. et al. De Novo Computational Design of Retro-Aldol Enzymes. Science 319, 1387–1391 (2008). 10.1126/science.1152692

13 Kim, D. et al. Computational design of metallohydrolases. Nature 649, 246–253 (2026). 10.1038/s41586-025-09746-w

14 Lauko, A. et al. Computational design of serine hydrolases. Science 388, eadu2454 (2025). 10.1126/science.adu2454

15 Röthlisberger, D. et al. Kemp elimination catalysts by computational enzyme design. Nature 453, 190–195 (2008). 10.1038/nature06879

16 Yeh, A. H.-W. et al. De novo design of luciferases using deep learning. Nature 614, 774–780 (2023). 10.1038/s41586-023-05696-3

17 Butcher, J. et al. De novo Design of All-atom Biomolecular Interactions with RFdiffusion3. bioRxiv (2025). 10.1101/2025.09.18.676967

18 Ahern, W. et al. Atom-level enzyme active site scaffolding using RFdiffusion2. Nat. Methods 23, 96–105 (2026). 10.1038/s41592-025-02975-x

19 Stark, H. et al. BoltzGen: Toward Universal Binder Design. bioRxiv (2025). 10.1101/2025.11.20.689494

20 Rector-Brooks, J. et al. General multimodal protein design enables dna-encoding of chemistry. arXiv preprint arXiv:2604.05181 (2026). 10.48550/arXiv.2604.05181

21 Cho, Y., Rangel, G., Bhardwaj, G. & Ovchinnikov, S. Protein Hunter: exploiting structure hallucination within diffusion for protein design. bioRxiv, 2025.10.10.681530 (2025). 10.1101/2025.10.10.681530

22 Fang, M. et al. HalluDesign: Protein Optimization and de novo Design via Iterative Structure Hallucination and Sequence design. bioRxiv, 2025.11.08.686881 (2025). 10.1101/2025.11.08.686881

23 Ross, S., Gordon, G. J. & Bagnell, J. A. A Reduction of Imitation Learning and Structured Prediction to No-Regret Online Learning. in Proceedings of the 14th International Conference on Artificial Intelligence and Statistics (AISTATS) (2011).

24 Bengio, S., Vinyals, O., Jaitly, N. & Shazeer, S. Scheduled Sampling for Sequence Prediction with Recurrent Neural Networks. in Advances in Neural Information Processing Systems (NeurIPS) (2015).

25. 25 Ho, J. & Salimans, T. Classifier-free diffusion guidance. arXiv preprint arXiv:2207.12598 (2022).

26. 26 Saharia, C., et al. Photorealistic Text-to-Image Diffusion Models with Deep Language Understanding. in Advances in Neural Information Processing Systems (NeurIPS) (2022).

27 Corley, N. et al. Accelerating Biomolecular Modeling with AtomWorks and RF3. bioRxiv, 2025.2008.2014.670328 (2025). 10.1101/2025.08.14.670328

28 Abramson, J. et al. Accurate structure prediction of biomolecular interactions with AlphaFold 3. Nature 630, 493–500 (2024). 10.1038/s41586-024-07487-w

29 Wohlwend, J. et al. Boltz-1: Democratizing Biomolecular Interaction Modeling. bioRxiv (2024). 10.1101/2024.11.19.624167

30 Dauparas, J. et al. Atomic context-conditioned protein sequence design using LigandMPNN. Nat. Methods 22, 717–723 (2025). 10.1038/s41592-025-02626-1

31 Rocha, J. F., Pina, A. F., Sousa, S. F. & Cerqueira, N. M. F. S. A. PLP-dependent enzymes as important biocatalysts for the pharmaceutical, chemical and food industries: a structural and mechanistic perspective. Catal. Sci. Technol. 9, 4864–4876 (2019). 10.1039/c9cy01210a

32 Tyka, M. D. et al. Alternate States of Proteins Revealed by Detailed Energy Landscape Mapping. J. Mol. Biol. 405, 607–618 (2011). 10.1016/j.jmb.2010.11.008

33 Beaudoin, S. F., Hanna, M. P., Ghiviriga, I. & Stewart, J. D. Progress in using threonine aldolases for preparative synthesis. Enzyme Microb. Technol. 119, 1–9 (2018). 10.1016/j.enzmictec.2018.07.004

34 Wang, J. et al. Scaffolding protein functional sites using deep learning. Science 377, 387–394 (2022). 10.1126/science.abn2100

35 Pacesa, M. et al. One-shot design of functional protein binders with BindCraft. Nature 646, 483– 492 (2025). 10.1038/s41586-025-09429-6

36 Tantillo, D. J., Chen, J. & Houk, K. N. Theozymes and compuzymes: theoretical models for biological catalysis. Curr. Opin. Chem. Biol. 2, 743–750 (1998). 10.1016/s1367-5931(98)80112-9

